# Repurposing ethacridine as a potent MMPL3 Inhibitor for the treatment of tuberculosis

**DOI:** 10.1101/2025.11.25.690360

**Authors:** Puja Kumari Agnivesh, Arnab Roy, Shashikanta Sau, Swati Meshram, Aman Dalal, Anamika Sharma, Nitin Pal Kalia

## Abstract

*Mycobacterium tuberculosis (Mtb),* the pathogen responsible for tuberculosis, remains a major global health threat, particularly with the rise of multidrug-resistant and extensively drug-resistant strains. This has renewed interest in repurposing existing drugs and exploring new cellular targets. The mycobacterial cell envelope is a key barrier to antibiotics and an attractive site for therapeutic intervention. In this study, we identify the FDA-approved drug ethacridine as a strong inhibitor of MmpL3, an essential transporter required for exporting trehalose monomycolate and building the cell wall. Computational docking and molecular dynamics indicate that ethacridine engages the MmpL3 binding pocket at residues also targeted by SQ109. Ethacridine shows potent activity against drug-sensitive and resistant *Mtb* isolates, with an MIC of 1 μg/mL, and remains effective against non-replicating bacteria and intracellular infection. Ethacridine-resistant mutants, overexpression strains, and a spheroplast TMM-flipping assay confirm MmpL3 as the target. The compound also disrupts the membrane potential, and flow-cytometry assays

## Introduction

*Mycobacterium tuberculosis (Mtb)* is one of the most ancient pathogens known to humans, responsible for tuberculosis (TB), a disease that continues to increase in prevalence. From 2021 to 2022, TB cases rose by 3.6%, with 10.1 million people infected in 2021 and 10.6 million cases reported in 2022. Between 2019 and 2021, TB-related deaths were substantial, with 1.4 million among HIV-negative individuals and 187,000 among HIV-positive individuals. The incidence of drug-resistant TB also worsened during this period (World Health Organization, 2022). Mycobacterial infections remain particularly difficult to treat due to the bacterium’s distinctive outer membrane, which is hydrophobic and limits the penetration of antibiotics. This unique structure not only helps the bacteria withstand hostile conditions but also diminishes the effectiveness of many conventional antibiotics. One family of proteins critical to the structure of the mycobacterial cell envelope is the mycobacterial membrane protein large (MmpL) family. Among these, Mycobacterial membrane protein large 3 (MmpL3) plays an essential role in transporting trehalose monomycolate (TMM) from the cytoplasm to the periplasm, which is vital for maintaining the integrity of the cell wall (Babii *et al*, 2024; Zhou *et al*, 2024). Inhibiting MmpL3 disrupts this process, compromising cell envelope formation and leading to bacterial death, making it a key target for TB drug development. MmpL3 a lipid transporter with 2,835 bp, 944 amino acid and 12 transmembrane (McNeil & Cook, 2019). It plays essential role in biosynthetic pathway of mycolic acid. It belongs to resistance-nodulation-division (RND) family of transporter. Initially MmpL3 act as multi-drug efflux pump in gram negative bacteria and lipid transporter in gram positive bacteria which helps in synthesis of multiple layers of cell envelope. In mycobacteria MmpL3 is mycolic acid flippase which transport intracellular trehalose monomycolate (TMM) to periplasm to form multilayer which affect the drug entry in bacteria and also neutralize host immune system (Adams *et al*). With the help of chaperone, TMM is transported across periplasm and serve as substrate for antigen 85 (Ag 85) complex for mycolyltransferases which further form mycolic acid chain from TMM to arabinogalactan and synthesize TDM. Apart from TMM transport, MmpL3 also transport lipid like phosphatidylethanolamine which provide rigidity and strength to cell wall (Xu *et al*, 2017b; CC *et al*, 2019). Many small molecules inhibitors are active against MmpL3 identify by high-throughput screen. Inhibition of MmpL3 render the transport of TMM from inner to outer membrane of cell resulting in TMM accumulation in the cytoplasm, and hampers TDM and mycolic acid formation, which weakens the cell wall and virulence of *Mtb* (Bhattarai *et al*, 2023). Over 20 chemical scaffolds have been developed targeting MmpL3, only SQ109 being the most advanced compound, having completed Phase IIb clinical trials (Li *et al*, 2014a).

Drug repurposing is a powerful strategy for identifying new TB treatments. This approach builds on the existing safety profiles and efficacy data of previously approved drugs, streamlining the development of novel therapies while reducing costs and accelerating timelines (Pinzi *et al*, 2024). Drug repurposing have been emerged as promising for treating TB. From oxazolidinone family, linezolid has shown good potential in patient suffering from MDR-TB. Similarly, clofazimine, is used for leprosy, is also effective against drug-resistant TB Notably, repurposed drugs like linezolid from the oxazolidinone class have shown promise in treating multidrug-resistant TB (MDR-TB). Similarly, clofazimine, originally developed for leprosy, has proven effective against drug-resistant forms of TB (Tovar-Nieto *et al*, 2024a). In this study, we explored the potential of the FDA-approved drug ethacridine (7-ethoxyacridine-3,9-diamine), known for its use as an antiseptic and abortifacient, in targeting MmpL3 in *Mtb*. Our aim is to evaluate its effectiveness as a novel approach to TB therapy.

## Results

### Screening of test molecules for antitubercular activity

The antitubercular efficacy of ethacridine was assessed using the microbroth dilution method in 96-well microtiter plates. This assay was conducted against *Mtb* strains mc^2^ 6230 and H37Ra, as well as *Mycobacterium bovis*, with the established MmpL3 inhibitor SQ109 included as a positive control. Ethacridine demonstrated notable inhibitory activity across all tested strains, exhibiting a minimum inhibitory concentration (MIC) of 1 μg/mL, comparable to that of SQ109 (Figure 1A) (Briffotaux *et al*, 2022).

**Figure 1:**
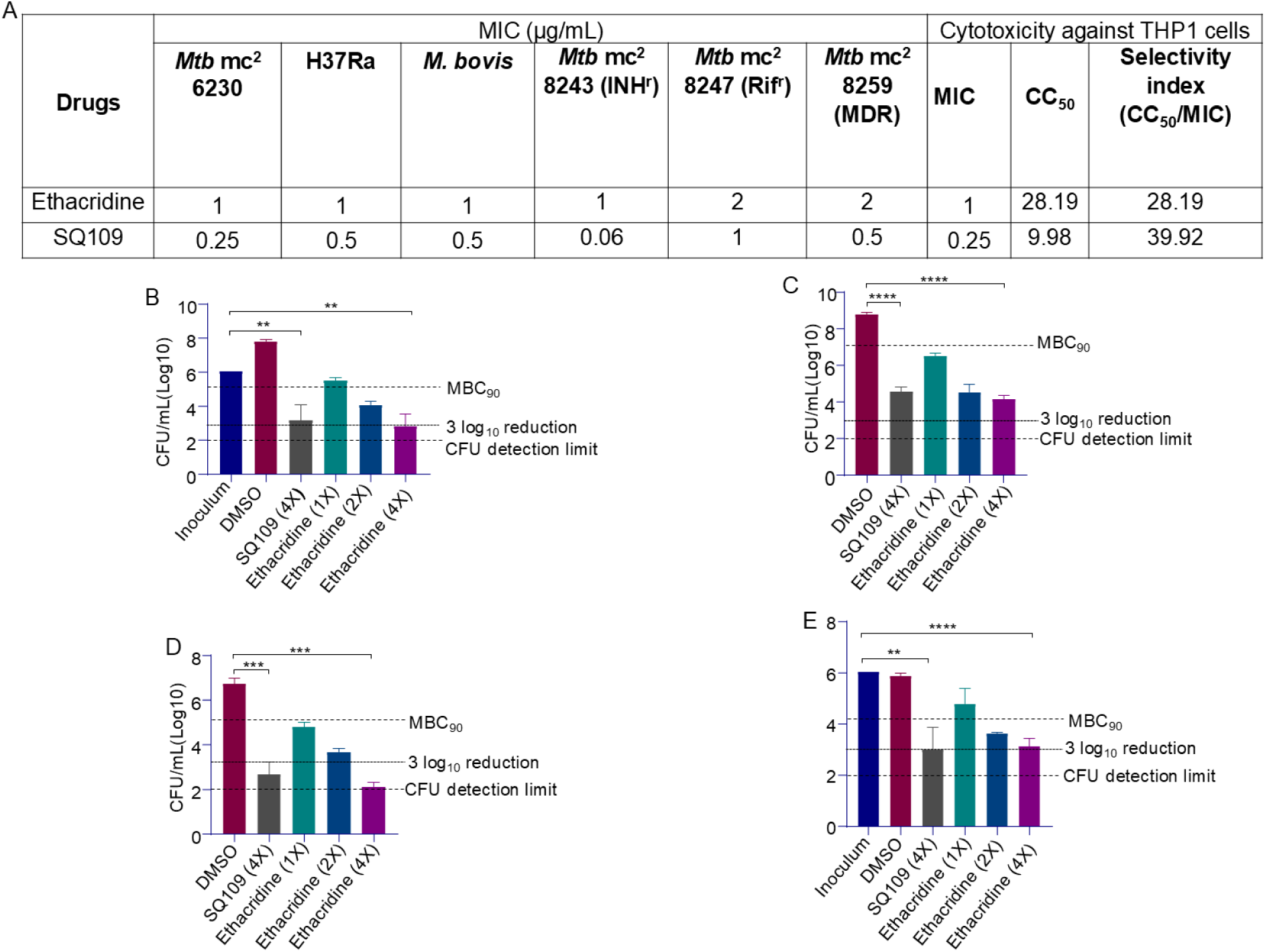
Potency of Ethacridine as an anti-TB agent: A) Minimum inhibitory concentrations (MICs) of ethacridine and SQ109 against various mycobacterial strains such as *Mtb* mc^2^ 6230, H37Ra, *M. bovis* as well as monoresistant strains, isoniazid-resistant (mc^2^ 8243), rifampicin-resistant (mc^2^ 8247), and multidrug resistant strain (mc^2^ 8259). Cytotoxicity of ethacridine and SQ109 was assessed in THP-1 cells using the MTT viability assay. Effect of ethacridine and SQ109 on the viability of *Mtb* mc^2^ 6230 after 10 days of treatment B) Minimum bactericidal concentration of ethacridine and SQ109 as control against susceptible *Mtb* mc^2^ 6230; C) Minimum bactericidal concentration of ethacridine and SQ109 as control against nutrient starved *Mtb* mc^2^ 6230; D) Minimum bactericidal concentration of ethacridine and SQ109 as control against hypoxia induced *Mtb* mc^2^ 6230; EA) Intracellular killing efficacy of ethacridine and SQ109 as control against *Mtb* H37Ra. Statistical significance was assessed using four-parameter logistic model to fit the log inhibitor concentration versus response curve, significance levels: p < 0.05 (*), p < 0.01 (**), p < 0.001 (***), and p < 0.0001 (****).

To further evaluate its therapeutic potential, ethacridine was tested against drug-resistant *Mtb* strains. In isoniazid-resistant (*Mtb* mc^2^ 8243) and rifampicin-resistant (*Mtb* mc^2^ 8247) strains, the compound displayed MIC values of 0.5 μg/mL and 1 μg/mL, respectively. For the multidrug-resistant strain *Mtb* mc^2^ 8259, the MIC remained at 1 μg/mL. In comparison, SQ109 exhibited MICs of 0.06 μg/mL (INH-resistant), 1 μg/mL (RIF-resistant), and 0.5 μg/mL (MDR strain) (Figure 1A). These findings indicate that ethacridine retains potent activity against both monoresistant and multidrug-resistant *Mtb*, supporting its potential utility as an anti-TB agent.

### Minimum Bactericidal Concentration of Ethacridine against distinct physiological states of *Mtb*

The minimum bactericidal concentration (MBC) of ethacridine was evaluated against *Mtb* mc^2^ 6230 under both replicating and non-replicating conditions, using SQ109 as a reference compound. Concentrations equivalent to 1x, 2x, and 4x the MIC were tested, with SQ109 at 4x MIC serving as the positive control. Under actively replicating conditions, ethacridine demonstrated a significant bactericidal effect, achieving a ≥3-log_10_ reduction in colony-forming units (CFU) at 4x MIC, comparable to the reduction observed with SQ109 (Figure 1B).

To assess activity against non-replicating bacilli, two *in vitro* models of dormancy were employed: hypoxia-induced and nutrient-starved cultures. In both models, ethacridine exhibited a dose-dependent reduction in bacterial viability. In nutrient-starved conditions, treatment with 4x MIC of ethacridine led to an approximate 3-log_10_ reduction in CFU, indicating strong bactericidal activity. A similar trend was observed in hypoxia-induced culture, where ethacridine also achieved a near-complete 3-log_10_ reduction in bacterial load at the same concentration (Figure 1B and 1C). These findings demonstrate that ethacridine retains potent bactericidal activity not only against actively growing *Mtb* but also against dormant bacilli, a major hurdle in current TB therapy (Figure 1C and 1D) (Li *et al*, 2014b).

### Cytotoxicity and Macrophage-Based Intracellular Activity of Ethacridine Against *Mtb*

We assessed the cytotoxicity of ethacridine and the reference drug SQ109 on THP1 cell lines using the MTT assay, which measures tetrazolium reduction. The half-maximal inhibitory concentrations (CC_50_) for both compounds were determined. Additionally, the selectivity index (SI) was calculated by dividing the CC_50_ by the MIC. Neither ethacridine nor SQ109 demonstrated significant cytotoxicity, with selectivity indices of 28.19 and 39.92 (Figure 1A), respectively. Compounds with an SI greater than 10 are generally considered to exhibit low toxicity and are deemed suitable for further investigation. Ethacridine’s ability to reduce the bacterial load increases as the concentration is raised. At 1x MIC, there is a modest reduction in bacterial numbers. As the concentration increases to 2x MIC, this reduction becomes more pronounced, indicating that ethacridine is effectively inhibiting bacterial count. At 4x MIC, ethacridine achieves a 3-log reduction in bacterial count (Figure 1E). This significant decrease suggests that ethacridine has intracellular killing efficacy. SQ109 reference for intracellular efficacy. Ethacridine’s 3-log_10_ reduction at 4x MIC is comparable to the results seen with SQ109, implying that ethacridine is just as effective in eliminating intracellular bacteria as SQ109. Ethacridine’s concentration-dependent efficacy and its ability to significantly reduce intracellular bacterial loads at higher concentrations suggest that it could be a strong candidate for treating tuberculosis.

### Susceptibility, Cross-resistance, and SNPs in MmpL3 gene

We investigated the potential of ethacridine as a MmpL3 inhibitor in lab-generated *Mtb* Ethacridine and SQ109 mutants. We selected random colonies for susceptibility testing after culturing them to the logarithmic growth phase. Among these, several cultures exhibited over an eight-fold increase in resistance, suggesting MmpL3 inhibition (Figure 2A). Cross-resistance assessments were conducted between ethacridine and SQ109-resistant mutants. Ethacridine displayed a four-fold increase in MIC against the SQ109-resistant strain, while SQ109 also showed an eight-fold increase in MIC when tested against the ethacridine-resistant strain, indicating possible overlapping binding sites. Subsequent sequencing of these mutants revealed point mutations in the MmpL3 gene, resulting in specific amino acid substitutions in the MmpL3 protein (Figure 2A). The frequency of spontaneous resistant mutants of *Mtb* was determined following exposure to ethacridine and SQ109 at 8x, 16x, and 32x to their respective MIC values. At 8x MIC, both compounds yielded mutant frequencies above 10^9^ CFU. Increasing the drug concentration to 16x MIC reduced the detectable mutant population, with ethacridine and SQ109 showing frequencies of 67 × 10^9^ and 97 × 10^9^ CFU, respectively. No resistant colonies were observed at 32x MIC for either compound, indicating frequencies below the detection limit (<10^9^ CFU) (Figure 2C)(Werngren, 2013). These observed mutations align with our *in-silico* binding predictions, where alterations occurred within the anticipated binding regions. Our findings support ethacridine’s role as an MmpL3 inhibitor, confirming its efficacy in targeting *Mtb* by disrupting MmpL3 function.

**Figure 2:**
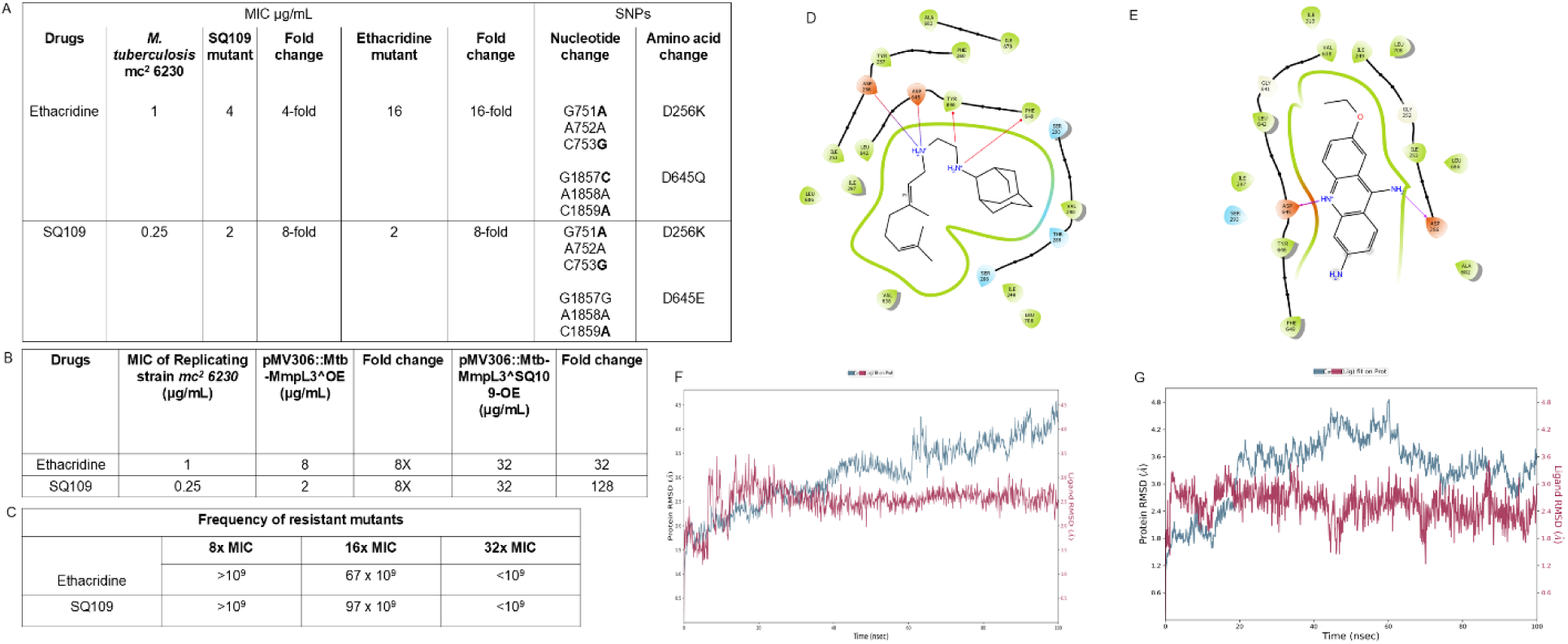
Ethacridine binds to MMPL-3 of *Mtb* and inhibit its function: A) SNP identification and cross-resistance testing between one step mutants of ethacridine and SQ109 demonstrated overlapping resistance profiles. B) Overexpression of MMPL-3 decreased the susceptibility of *Mtb* by 8–32-fold increase in MIC values consistent with target-associated resistance. C) Frequency of mutant generation D, E) Two-dimensional ligand interaction diagrams showing SQ109 D) and ethacridine E) binding within the MmpL3 active site, highlighting key hydrogen bonds and hydrophobic contacts. F, G) Molecular dynamics simulation trajectories depicting the RMSD profiles of the protein-ligand complexes over 100 ns for SQ109 F) and ethacridine G), indicating overall conformational stability during the simulation.

### Target Confirmation of Ethacridine through MmpL3 Overexpression in *Mtb*

In this study, we investigated the interaction of ethacridine with MmpL3, an essential transporter involved in mycolic acid translocation across the *Mtb* inner membrane and a validated target for anti-TB drug development. To evaluate the mechanism of action of ethacridine, we constructed *Mtb* strains overexpressing wild-type MmpL3 (pMV306::*Mtb*-MmpL3^OE) and SQ109-resistant variant of MmpL3 (pMV306::*Mtb*-MmpL3^SQ109-OE) using the integrative pMV306 plasmid. These constructs were transformed into *Mtb* mc^2^ 6230, and overexpression was verified through drug susceptibility testing. MICs of ethacridine and the reference MmpL3 inhibitor SQ109 were determined using a microbroth dilution assay. In the wild-type strain, the MICs for ethacridine and SQ109 were 1 µg/mL and 0.25 µg/mL, respectively. In contrast, the MmpL3 overexpression strain exhibited significantly elevated MIC of 8 µg/mL for ethacridine and 2 µg/mL for SQ109 (Figure 2B). Furthermore, the strain expressing the SQ109-resistant MmpL3 mutant displayed even higher resistance, with an MIC of 32 µg/mL for ethacridine.

The observed 8 to 32 fold increase in MICs in the overexpression and mutant strains strongly suggests a target-based mechanism, where elevated levels of MmpL3 or resistance-associated mutations diminish the efficacy of ethacridine.

### MmpL3 Mediates TMM Translocation Across the Inner Membrane

MmpL3 is implicated in the translocation of trehalose monomycolates (TMMs) across the inner membrane (IM). Small molecules such as SQ109, BM212, and AU1235 are known to interfere with this process, as resistance-conferring mutations frequently map to the *mmpL3* gene. To assess the potential of ethacridine as a TMM flippase inhibitor, spheroplast-TMM flippase assay were performed using concentrations equivalent to the MIC and 4x MIC. SQ109 at 4x MIC was used as a positive control. Spheroplasts treated with DMSO showed strong fluorescence, indicating efficient translocation of newly synthesized TMM to the outer leaflet of the inner membrane. In contrast, exposure to ethacridine led to a dose-dependent reduction in fluorescence intensity, with the 1x MIC and 4x MIC treatments showing substantial decreases compared to the control. SQ109, used as a reference MmpL3 inhibitor, produced a similar reduction, though the magnitude was lower than that observed with high-dose ethacridine (Figure 3B). Quantification of the fluorescence signal revealed inhibition levels of 54.8% and 67.4% for ethacridine at 1x and 4x MIC, respectively, and 43.1% for SQ109 (Figure 3C). Fluorescence microscopy corroborated these findings, DMSO-treated spheroplasts exhibited bright surface labeling, whereas cells treated with ethacridine or SQ109 showed markedly diminished signal (Figure 3D). These findings support the conclusion that ethacridine inhibits MmpL3-mediated TMM flipping, thereby validating it as an MmpL3-targeting compound.

**Figure 3:**
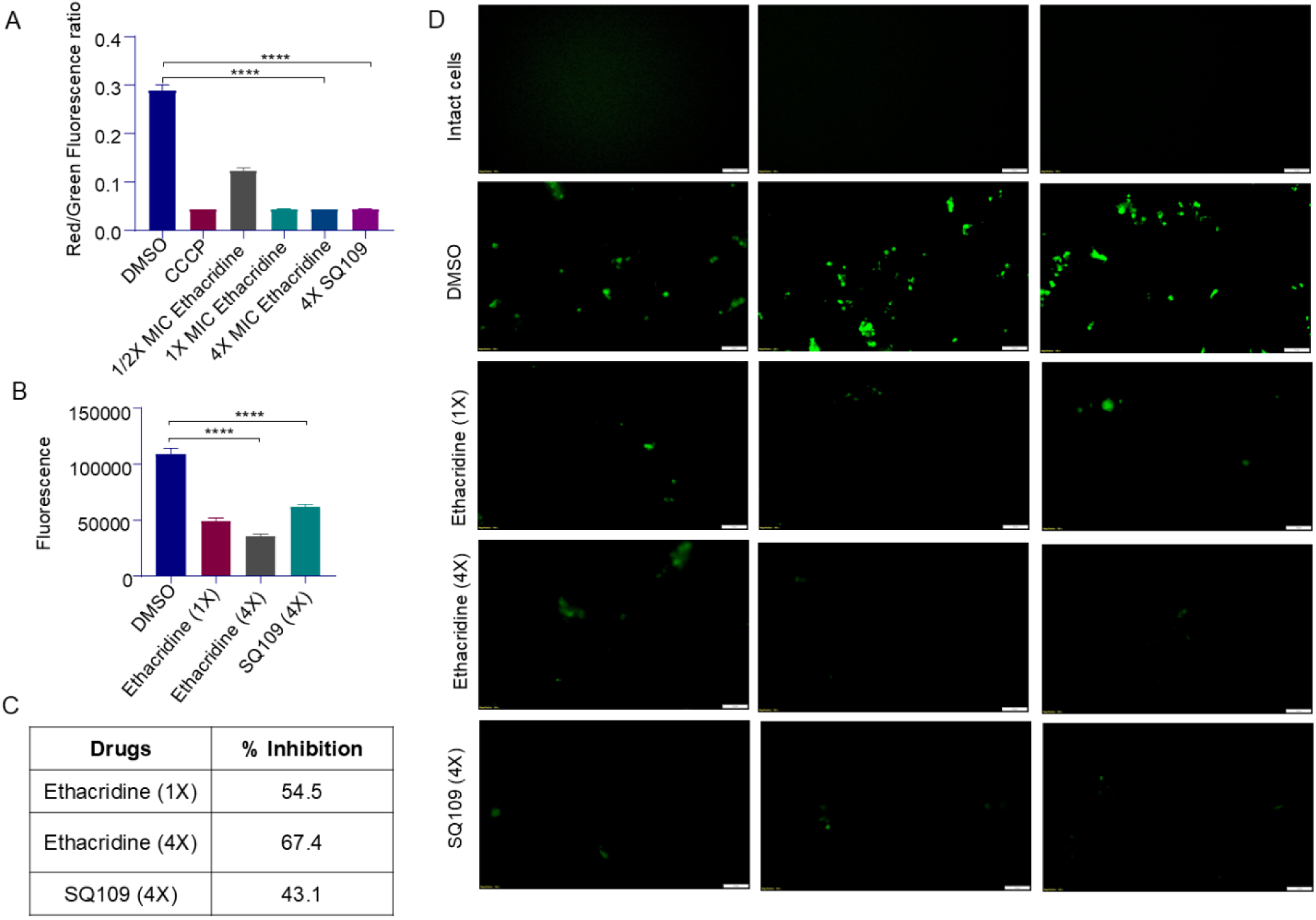
Effect of Ethacridine on membrane potential and MmpL3 dependent TMM translocation: A) Membrane potential (ΔΨ) was measured using DiOC₂ dye. Ethacridine and SQ109 were tested at concentrations of ½x, 1x, and 4x MIC. Carbonyl cyanide m-chlorophenyl hydrazone (CCCP) at 25 μM served as a standard protonophore, while DMSO was used as a control at a concentration equivalent to the maximum amount used in treatments (kept below 1%). B) Inhibition of MmpL3-mediated translocation of 6-azido-TMM in spheroplasts following treatment with ethacridine (1x and x MIC) or SQ109 (4x MIC). Fluorescence corresponding to surface-exposed TMM (Ex 488 nm/Em 525 nm) was quantified relative to the DMSO control. C) Percentage inhibition of TMM flipping calculated from fluorescence measurements shown in panel B. D) Representative fluorescence microscopy images of spheroplasts labeled with 6-azido-TMM. Untreated intact cells show negligible signal, while DMSO-treated spheroplasts display strong surface fluorescence. Treatment with ethacridine or SQ109 results in markedly reduced fluorescence, consistent with impaired TMM translocation across the inner membrane. Scale bars correspond to the magnification used in each panel.

### Evaluation of Ethacridine-Induced Changes in Mycobacterial Membrane Potential (ΔΨ) via DiOC_₂_

The membrane potential (ΔΨ) plays a crucial role in maintaining the energy homeostasis of *Mtb*, and disruption of ΔΨ is a promising strategy for combating this pathogen. In this study, we investigated the effects of MmpL3 inhibitors, SQ109, and ethacridine, on the ΔΨ of *Mtb*. To assess membrane depolarization, we utilized CCCP as a standard protonophore and DMSO as a negative control. The ΔΨ was quantified using red/green fluorescence measurements, providing insights into membrane integrity and potential disruptions caused by the inhibitors. We administered ethacridine and SQ109 at concentrations of 1/2x MIC, 1x MIC and 4x MIC to evaluate concentration-dependent effects on ΔΨ. At sub-MIC levels (1/2x MIC), both ethacridine and SQ109 exhibited a modest reduction in the red/green fluorescence ratio, suggesting an initial disruption of the ΔΨ. However, at 1x MIC and 4x MIC, the decrease in fluorescence was pronounced, reaching levels comparable to that induced by CCCP (Figure 3A). This significant reduction in fluorescence indicates substantial membrane depolarization, underscoring the capacity of ethacridine and SQ109 to disrupt membrane integrity at elevated concentrations. Our findings reveal that both ethacridine and SQ109 effectively disrupt the ΔΨ of *Mtb* at higher concentrations, paralleling the action of standard protonophores. These results underscore the potential of MmpL3 inhibitors as therapeutic agents targeting the bacterial membrane, thereby contributing to *Mtb* eradication through ΔΨ disruption.

### Flow Cytometry-Based Viability Analysis Using Sytox Green and Propidium Iodide

Drug treatment caused a dose-dependent increase in cells with compromised membranes, as evidenced by elevated fluorescence in SYTOX Green (SG) and Propidium Iodide (PI) assays. In SG-stained samples (Figure 4C), control cultures showed minimal fluorescence, indicating intact membranes and low background cell death. Ethacridine at 4x minimum inhibitory concentration (MIC) resulted in 35.85 ± 3.95% SG-positive cells, while SQ109 at 4x MIC led to 61.31 ± 3.57% membrane-compromised cells. A positive control exhibited 98.95 ± 0.66% SG-positive cells, confirming SG’s efficacy in detecting cell death.

**Figure 4:**
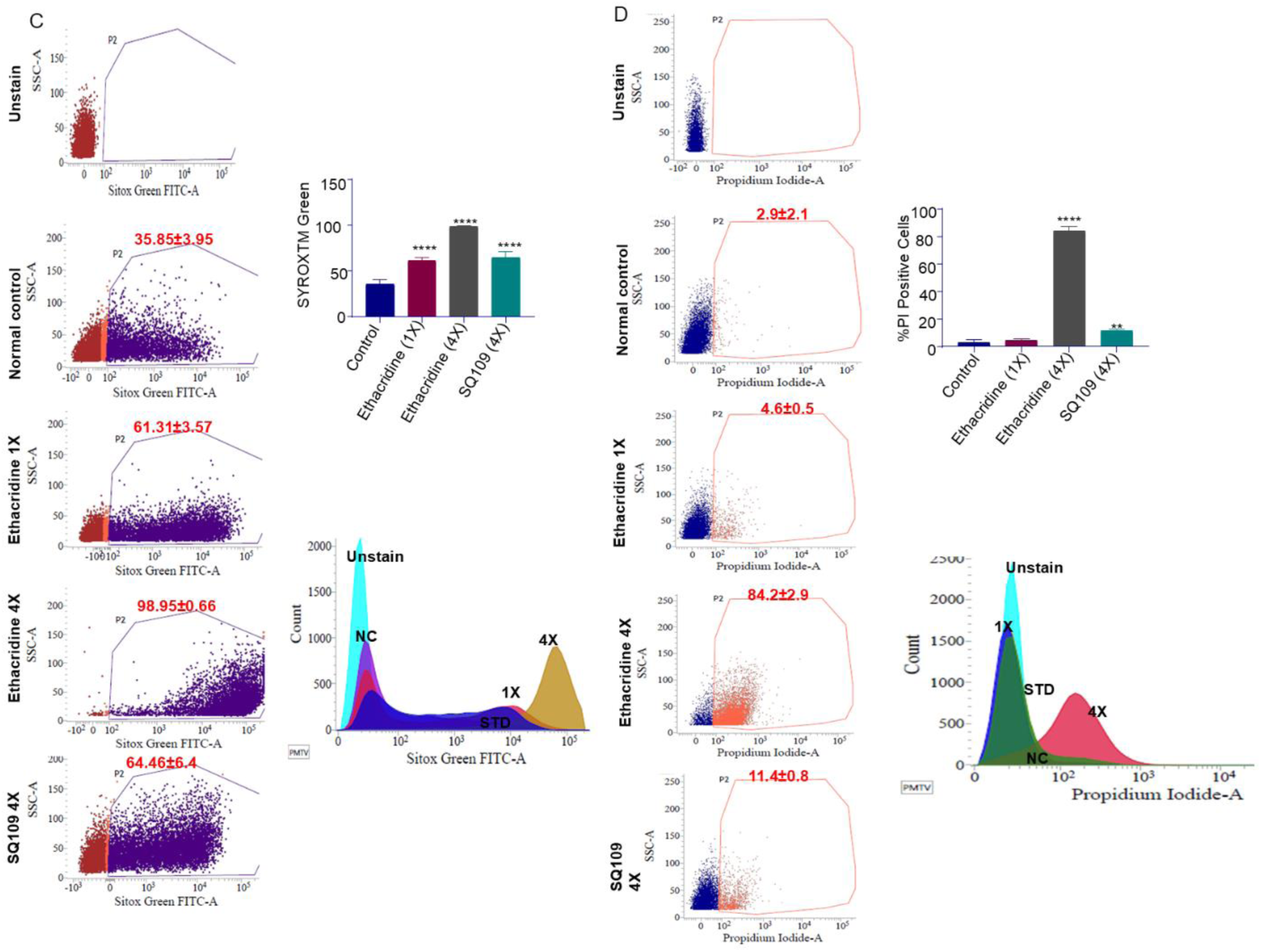
Ethacridine-induced membrane depolarization and permeability changes in *Mycobacterium tuberculosis* assessed by flow cytometry: C and D) Flow cytometric analysis of *Mycobacterium tuberculosis* viability using Sytox Green and Propidium Iodide staining. Bacterial cultures were treated with indicated concentrations (4× MIC) of Ethacridine and SQ109, followed by staining with Sytox Green (SG) or Propidium Iodide (PI) to assess membrane integrity. C) Dot plots and corresponding histograms show SG fluorescence intensity in untreated (NC), Ethacridine-treated, and SQ109-treated populations. A shift in fluorescence toward the right (P3 gate) indicates an increase in non-viable, membrane-compromised cells. Quantitative analysis revealed 35.85 ± 3.95% SG-positive cells in Ethacridine-treated samples and 61.31 ± 3.57% in SQ109-treated cells. A positive control demonstrated near-complete staining (98.95 ± 0.66%). D) PI-based analysis showing similar dot plots and histograms. Drug-treated samples exhibited increased PI uptake, confirming compromised membrane integrity. SQ109 treatment produced the highest shift in fluorescence intensity, consistent with its stronger bactericidal activity. Overlay histograms depict SG and PI signal intensities in unstained, 1X, and 4X MIC-treated groups. Increased fluorescence with drug concentration further supports dose-dependent loss of bacterial viability.Statistical significance was assessed using four-parameter logistic model to fit the log inhibitor concentration versus response curve, significance levels: p < 0.05 (*), p < 0.01 (**), p < 0.001 (***), and p < 0.0001 (****).

Similarly, PI-stained samples (Figure 4D) corroborated the SG findings. Untreated cells displayed low PI fluorescence, consistent with a predominantly viable population. Ethacridine-treated samples showed increased PI-positive cells, while SQ109 treatment resulted in a more pronounced rightward shift in fluorescence intensity, indicating greater membrane damage. Histogram overlays and population distribution analyses confirmed that ethacridine and SQ109 exerted potent bactericidal effects.

Overlay histograms comparing unstained, 1x MIC, and 4x MIC drug-treated populations highlighted drug-induced increases in fluorescence intensity. Flow cytometry data from both SG and PI assays demonstrate that these compounds disrupt bacterial membrane integrity, directly contributing to reduced cell viability.

### Molecular Docking and Molecular Dynamics Simulation Studies of Ethacridine

The compelling antitubercular efficacy of ethacridine led us to explore its interaction with the binding site of the MmpL3 protein using molecular docking and molecular dynamics (MD) simulation studies. Given the limited structural data for *Mtb* MmpL3, we employed the X-ray crystal structure of MmpL3 from *Mycobacterium smegmatis* (PDB: 6AJH) due to its sequence similarity with *M tuberculosis*. Docking analysis indicated that ethacridine occupies the same binding pocket as SQ109, an established MmpL3 inhibitor (Figure 2D and E). The amide NH group of ethacridine demonstrated critical hydrogen bonding with Asp645 and formed two aromatic hydrogen bonds with Asp645 and Asp256, which stabilized the binding pose, closely resembling that of SQ109. A 100ns MD simulation was conducted on the protein-ligand complex to evaluate any structural modifications upon ligand binding. Root mean square deviation (RMSD) analyses of the protein and ligand were performed for both SQ109 (as the reference) and ethacridine to assess the stability of the complex. The RMSD values for ethacridine and SQ109 with respect to the protein remained consistently below 4.5 Å, indicating stable binding throughout the simulation (figure 2F and G). This study underscores the significance of ethacridine in targeting MmpL3 and encourages further investigation into its potential as an antitubercular agent.

## Discussion

The cell wall of *Mtb* is a complex, multi-layered structure that includes the outer mycomembrane, acting as a formidable barrier that limits antibiotic penetration and shields the bacteria from host immune responses. This cell envelope significantly contributes to drug resistance by reducing antibiotic access. A major hurdle in TB treatment is achieving complete sterilization of *Mtb*, as partial clearance can promote the rise of drug-resistant strains and lead to infection relapse. One promising therapeutic target is the mycobacterial membrane protein large (MmpL) family, particularly MmpL3, which is essential for the synthesis and transport of complex cell envelope lipids. MmpL proteins are crucial for lipid transport across the membrane, as confirmed by genome sequencing, with MmpL3 playing a key role in bacterial viability. Inhibiting MmpL3 disrupts cell wall synthesis, which leads to bacterial death, making it a focal point for TB drug development (Bolla, 2020; Melly *et al*, 2019). *Mtb* has 14 MmpLs which helps in lipid transportation confirmed by genome sequencing. Out of all membrane transport protein only MmpL3 is essential for maintaining the viability of bacteria as it is a key for the synthesis of complex envelope and inhibiting MmpL3 activity will leads to arrest of cell wall synthesis and ultimately cell death (Delmar *et al*, 2015; Bolla, 2020) Among cell wall synthesis inhibitors, SQ109, an MmpL3 inhibitor, is currently in Phase IIb/III clinical trials (Li *et al*, 2023). In light of the lengthy timelines and high costs associated with traditional drug discovery, drug repurposing offers an efficient alternative by leveraging the established safety, pharmacokinetic, and toxicological profiles of existing drugs, as illustrated by the use of linezolid in TB treatment. Integrating both novel and repurposed antimycobacterial is critical to addressing resistance, shortening treatment duration, and minimizing toxicity, which aligns with the goals of the End TB Strategy (Hanscheid *et al*, 2024; Sharma *et al*, 2023). In our study, we explored the FDA-approved drug ethacridine, known for its antiseptic properties and medical abortion, as a potential anti-tubercular agent. Ethacridine showed efficacy across several Mycobacterial strains, including *Mtb* mc^2^ 6230, H37Ra, and *M. bovis*, with a MIC of 1 µg/mL and MIC_50_ of <0.5 µg/mL, which was comparable to the reference drug SQ109. The drug exhibited bactericidal activity and a time-dependent killing effect. In nutrient-starved and drug-susceptible *Mtb* mc^2^ 6230 strains, ethacridine demonstrated a 3-log reduction at 4x MIC, reducing bacterial load to below the colony-forming unit (CFU) detection limit.

Ethacridine was also effective against drug-resistant strains, including isoniazid-resistant (mc^2^ 8243) and rifampicin-resistant (mc^2^ 8247) strains, with MIC values of 0.5 and 1 µg/mL, respectively. In multi-drug-resistant strain (mc^2^ 8259), it maintained an MIC of 1 µg/mL, highlighting its broad-spectrum anti-tubercular potential. Cytotoxicity assays using the THP-1 cell line indicated a selectivity index >10, confirming ethacridine’s safety profile. Furthermore, intracellular killing assays demonstrated activity at 2x and 4x MIC, and at 4x MIC activity is comparable to SQ109.

To identify and validate the molecular target of ethacridine, single-step spontaneous resistant mutants were isolated by exposing Mycobacterium tuberculosis to ethacridine and the reference compound SQ109. Mutants selected independently for each compound exhibited cross-resistance, indicating a likely shared mechanism of action. Whole-genome sequencing of the resistant clones revealed point mutations localized to the mmpl3 gene, a known target of SQ109. These findings were consistent with molecular docking simulations, which predicted that both compounds bind to overlapping sites within the transmembrane region of the MmpL3 transporter, suggesting a common binding pocket.

To further validate the target, *Mtb* strains overexpressing MmpL3 were constructed. These strains displayed an 8- to 32-fold elevation in the MIC for ethacridine, confirming that increased MmpL3 levels reduce drug susceptibility and supporting MmpL3 as the primary target of action. Functional validation was carried out using a spheroplast-based trehalose monomycolate (TMM) flippase assay, which showed that ethacridine impairs TMM translocation, a process mediated by MmpL3, thereby disrupting mycolic acid transport and cell wall biosynthesis. Additionally, the effect of ethacridine on the proton motive force (PMF) was assessed. Treatment with ethacridine led to a significant disruption of the transmembrane potential (ΔΨ), a key component of PMF, further supporting its action on membrane-associated targets. Flow cytometric analysis using membrane-impermeable dyes such as Sytox Green and Propidium Iodide confirmed ethacridine-induced membrane damage, consistent with its mechanism of action involving MmpL3 inhibition and disruption of cell envelope integrity.

## Materials and Methods

| Reagents and tool table |  |  |
| --- | --- | --- |
| Reagent/Resource | Reference or Source | Identifier or Catalog Number |
| **Experimental Models** |  |  |
| *Mycobacterium bovis* (*M. bovis*) | Laboratory stock |  |
| H37Ra | ATCC, USA |  |
| *Mtb* mc<sup>2</sup> 6230 | Prof. William Jacob Jr.,<br>Albert Einstein College of<br>Medicine, USA. |  |
| *Mtb* mc<sup>2</sup> 8243 |  |  |
| *Mtb* mc<sup>2</sup> 8247 |  |  |
| *Mtb* mc<sup>2</sup> 8259 |  |  |
| DH5α | Laboratory stock | |
| pMV306 | Addgene | 26155 |
| THP1 | ATCC | ATCC#TIB-202 |
| **Oligonucleotides and other sequence-based** |  |  |

| reagents |  |  |
| --- | --- | --- |
| Primers | Eurofins |  |
| EcoRI | NEB | R0101S |
| HIND III | NEB | R0104S |
| DNA Ladder | NEB | N3232L |
| GoTaq Master Mixes | Promega | M7132 |
| Phorbol-12-myristate 13-Acetate | Krishgen Biosystem | 10008014 |
| Chemicals, Enzymes and other reagents |  |  |
| Middlebrook 7H9 Broth Base | BD Difco Laboratories, USA | 271310 |
| Middlebrook 7H10 Agar | BD Difco Laboratories, USA | 262710 |
| Agarose | Sigma-Aldrich | A4718 |
| Bovine serum albumin | Sigma-Aldrich | TC194 |
| DMSO | Sigma-Aldrich | 472301 |
| SQ109 | Sigma-Aldrich | G2964-85005 |
| Tween 80 | Sigma-Aldrich | P4780 |
| L-leucine | Sigma-Aldrich | L8000 |
| L-Glutamine | Sigma-Aldrich | MV1245 |
| Beta mercapto ethanol | Sigma-Aldrich |  |
| MTT | Sigma-Aldrich | M5655 |
| L-methionine | Sigma-Aldrich | 1057070025 |
| RPMI-1640 | Sigma-Aldrich | R0883-6 |
| Glycerol | Sigma-Aldrich | G5516 |
| Carbonyl Cyanide m-Chlorophenylhydrazone | Sigma-Aldrich | C1016 |
| Calcium Chloride | Sigma-Aldrich | C1016 |
| 2-Mercapto Ethanol | Sigma-Aldrich | M6250 |
| EDTA | Sigma-Aldrich | E9884 |
| L-Glutamine | Sigma-Aldrich | G7513 |
| SDS | Sigma-Aldrich | 74255 |
| Luria Bertani Broth | Himedia | MV1245 |
| Kanamycin | Himedia | TC136M |
| Propidium Iodide | Himedia | MB139 |
| Isoniazid | Himedia | CMS7169 |
| Rifampicin | Himedia | TC354 |
| Lysozyme | Himedia | MB098 |
| Streptomycin | Himedia | 27958 |
| Penicillin G | Himedia | TC187 |
| Magnesium sulphate | Himedia | MB171 |
| Sucrose | Himedia | MB025 |
| Ferric Ammonium Sulphate | Himedia | PCT0107 |
| Catalase | Himedia | TC037 |
| Quick CIP | NEB | M0525S |
| Ethidium Bromide | NEB | MB074 |
| SYTOX Green | Invitrogen | S7020 |
| Streptavidin-Alexa Fluor | Invitrogen | S11223 |
| Click-iT™ sDIBO Alkynes for copper-free click chemistry | Invitrogen | C20023 |
| DIOC<sub>2</sub> | Invitrogen | D14730 |
| DPBS | Invitrogen | 14200075 |
| D-pantothenate | SRL | 31269 |
| L-arginine | SRL | 66637 |
| 6-Azido trehalose | Jena Biosciences | CLK-078 |
| FBS | Thermofisher Scientific | 26140079 |
| HEPES Buffer | Gibco | 2193374 |
| **Software** |  |  |
| GraphPad Prism8 | Institute subscription |  |
| Maestro software's 14.4 | Institute subscription |  |
| **Other** |  |  |
| Whole Genome Sequencing | Eurofin Genomics India Private Limited |  |
| Anaerobic atmosphere generation bags | Sigma-Aldrich | 68061 |
| Gel Purification Kit | ILS | G214201921 |
| PCR Cleanup Kit | Sigma-Aldrich | NA1020-1KT |
| Plasmid Isolation Kit | Invitrogen | K210010 |
| Quick Ligation Kit | NEB | M2200S |

### Bacterial strains, cell lines, drugs, and media

For cloning *Escherichia coli* strain DH5α were used, it was grown in LB broth (Himedia). For evaluation of antitubercular activity bacterial strain such as H37Ra ATCC25177, *Mycobacterium bovis,* and auxotrophic *Mtb* mc^2^ 6230 and its derivative strains were used. These strains were grown in 7H9 liquid broth and 7H10 solid agar which were supplemented with oleic-albumin-dextrose-catalase (OADC), 0.2% glycerol, and 0.05% Tween 80 (BD Difco Laboratories, USA). Auxotrophic strains were grown with additional nutrients supplementation such as L-leucine (50 mg/L), D-pantothenate (24 mg/L), L-methionine (50 mg/L), and L-arginine (Sigma-Aldrich). Non-replicating nutrient starvation culture were generated in DPBS supplemented with calcium and magnesium (Sigma-Aldrich). The cell line THP1(ATCC#TIB-202) were grown RPMI media supplemented with 10% fetal bovine serum (FBS) (Gibco). The anti-TB drugs such as isoniazid, rifampicin, and SQ109 were purchased from Sigma Aldrich. Drugs were prepared at a concentration of 10mg/mL in DMSO purchase from Sigma Aldrich.

### Determination of Minimum inhibitory concentration and Minimum bactericidal concentration

The antitubercular efficacy of the test compound was assessed using a microbroth dilution technique in 96-well microtiter plates against various Mycobacterium strains. The bacterial cultures were grown to the logarithmic phase (OD_600_ between 0.4 and 0.6) in 7H9 broth supplemented ADC. For MIC determination, the cultures were seeded at OD_600_ of 0.005 (approximately 2 x 10^5^ CFU/mL) in 96-well round-bottom plates, while MIC_50_ assays were conducted in flat-bottom plates. Both assays involved testing the compound and standard antimycobacterial drugs in two-fold serial dilutions, with a final volume of 200 μL per well. The plates were then incubated at 37°C for 10 days. MIC values were determined visually based on turbidity (Shetty *et al*, 2018b; Foss *et al*, 2016a).

To determine the minimum bactericidal concentration (MBC) of the test compound, assays were conducted using *Mtb* mc^2^ 6230 under three physiological states: replicating, hypoxia-induced non-replicating, and nutrient-starved non-replicating conditions. For the replicating condition, *Mtb* mc^2^ 6230 was cultured to mid-log phase (OD_600_: 0.4–0.6) and subsequently diluted to an OD_600_ of 0.005 for the assay setup. To simulate hypoxic non-replicating conditions, cultures were adjusted to an OD_600_ of 0.1 and incubated in sealed tubes with anaerobic sachets. Methylene blue served as an oxygen indicator, where decolorization indicated the establishment of hypoxic conditions, as previously described by Kalia et al (Kalia *et al*, 2023) For nutrient starvation-induced dormancy, bacilli in exponential phase (OD_600_: 0.1–0.2) were harvested by centrifugation, washed with phosphate-buffered saline (PBS) supplemented with calcium, magnesium, and 0.025% Tween 80, and resuspended in the same buffer to an OD_600_ of 0.15. Cultures were then incubated under starvation conditions for 14 days following the protocol described by Gengenbacher et al (Gengenbacher *et al*, 2010). MBC assays were performed by testing the drug at concentrations equivalent to 1x MIC, 2x MIC, and 4x MIC, with SQ109 at 4x MIC used as a positive control, and DMSO as the negative control. Aliquots from each serial dilution were plated onto agar and incubated at 37°C for 3–4 weeks. Following incubation, the colonies were counted, and the colony-forming units (CFU) per mL were calculated to evaluate the bactericidal activity of the test compound (Shetty *et al*, 2018b).

### Activity against resistant strains

Ethacridine was evaluated against both mono drug-resistant and multidrug-resistant mycobacterium strains using the microbroth dilution method. The monodrug-resistant strains included an isoniazid (INH^r^)-resistant strain (*Mtb* mc^2^ 8243) and a rifampicin (RIF^r^)-resistant strain (*Mtb* mc^2^ 8247), alongside a multidrug-resistant strain resistant to both INH and RIF (*Mtb* mc^2^ 8259).

### Cytotoxicity assay

The cytotoxicity of the ethacridine was evaluated using THP1 human monocytic cell lines. Cells were cultured in RPMI media supplemented with 10% heat inactivated fetal bovine serum (FBS) (Gibco) to approximately 80% confluency before being seeded into 96-well, tissue culture-treated microtiter plates at a density of 50,000 cells per well with PMA (20 ng/mL) for 48 hours. Serial two-fold dilutions were prepared, starting at an initial concentration of 256 µg/mL, with tamoxifen as the positive control. After a 24-hour incubation at 37°C in a 5% CO₂ atmosphere, the media was replaced with fresh media containing 2mg/mL MTT [3-(4,5-dimethylthiazol-2-yl)-5-(3-carboxymethoxyphenyl)-2-(4-sulfophenyl)-2-tetrazolium inner salt]. The plates were incubated for another 2 hours under the same conditions. Following incubation, the media was aspirated, and the resulting formazan crystals were dissolved in DMSO. Absorbance was measured at 570 nm using a BioTek Citation 5 multimode reader. CC_50_ values were calculated with GraphPad Prism 5 software, and the selectivity index (SI) was derived by calculating the ratio of CC_50_ to MIC (Shetty *et al*, 2018c) (Foss *et al*, 2016b).

### Intracellular assay

The THP-1 cell line was cultured in RPMI 1640 medium, and seeded (2 x 10^5^ cells/well) and infected with *Mtb* H37Ra strain (2 x 10^6^) at a multiplicity of infection (MOI) of 10 bacilli per cell. After 4 hours of incubation to allow infection, the cells were washed thoroughly with antibiotic-free RPMI 1640 medium to remove any extracellular bacilli, ensuring only intracellular bacteria remained. Following infection, the THP-1 cells were treated with ethacridine at concentrations of 1x, 2x, and 4x MIC. SQ109 at 4x MIC was used as a positive control, and DMSO-treated cells served as a negative control. All treatments were performed in triplicate. The cells were incubated with the respective compounds for a period of 5 days. After the treatment period, infected THP-1 cells were lysed using 0.01% SDS, a mild detergent, to selectively lyse the host cells and release the intracellular *Mtb*. The lysates were serially diluted and plated onto Middlebrook 7H10 agar. The plates were incubated at 37°C for 3 weeks to allow the formation of visible colonies. Colony-forming units (CFU) were counted to assess the number of viable *Mtb* bacilli following the treatments (Tovar-Nieto *et al*, 2024b).

### Target confirmation through one-step mutant generation

To investigate whether ethacridine shares a binding site with the known MmpL3 inhibitor SQ109, we generated single-step mutants resistant to each compound. Specifically, *Mtb* mc^2^ 6230 was exposed to 8x, 16x and 32x MIC concentrations of SQ109 and ethacridine on 7H10 agar plates supplemented with 10% OADC. A high bacterial inoculum (10^9^ CFU/mL) was applied to ensure mutation selection. Plates were incubated at 37°C for three weeks, after which resistant colonies were isolated and propagated in 7H9 broth supplemented with 10% ADC until reaching logarithmic growth. To assess potential cross-resistance, the mutant’s susceptibility to both SQ109 and ethacridine was determined using a microbroth dilution assay. Mutants exhibiting resistance levels exceeding eight times the MIC for both compounds, as compared to the wild-type strain, were further examined. The MmpL3 encoding gene of these high-level resistant mutants was sequenced to detect single nucleotide polymorphisms (SNPs), providing insights into the binding interactions and potential shared target sites of SQ109 and ethacridine (Williams *et al*, 2019)(Werngren, 2013).

### Overexpression of MmpL3 through pMV306

The MmpL3 gene region (2835 bp) was amplified using polymerase chain reaction (PCR) from genomic DNA isolated from *Mtb* mc^2^ 6230 and SQ109 mutant. Primers used for this amplification were 5’-TTGCCTGAATTCGTGTTCGCCTGGTGGGGTCG-3’ and 5’-TCTCGGCAAGCTTTTAAAGGCGTCCTTCGCGGC-3’. The amplified product was cloned into the pMV306 vector under the control of the hsp promoter. Both the amplified MmpL3 fragment and the pMV306 vector were purified and digested with EcoRI and HindIII restriction enzymes. Following digestion, the products were run on 0.8% agarose gel and subsequently purified using the Invitrogen Quick Gel Extraction Kit (Shetty *et al*, 2018a)(Stover *et al*, 1991). The digested MmpL3 fragment was ligated into the vector using Quick Ligase (Invitrogen) and transformed into *Escherichia coli* DH5α by the heat shock method. Transformed colonies were plated on LB agar supplemented with 50 µg/mL kanamycin to select for positive constructs. Colonies were screened for the presence of the MmpL3 construct through restriction enzyme digestion. The confirmed construct was then electroporated into *Mtb* mc^2^ 6230 using a Bio-Rad Gene Pulser set to 2500 V, 1000 Ω, and 25 µF, with 2-mm cuvettes. Cells were initially grown overnight in 7H9 broth before being plated onto 7H10 agar containing 20 µg/mL kanamycin. (He *et al*, 2024). Plates were incubated at 37°C until colonies appeared. Individual colonies were selected and checked for MmpL3 overexpression by microbroth dilution. Overexpression was inferred by an increased minimum inhibitory concentration (MIC) of drugs compared to the wild-type strain, consistent with reduced drug susceptibility due to MmpL3 overexpression.

### Spheroplast-Based TMM flippase assay

Spheroplasts of *Mtb* mc² 6230 were generated following the method described by Thacore and Willett (1963) (Thacore & Willett, 1963). These spheroplasts were metabolically labeled with 6-azido-trehalose for 2 hours at 37 °C to facilitate the biosynthesis of 6-azido-trehalose monomycolate (6-azido-TMM). Subsequently, a copper-free click reaction was performed using Click-IT™ Biotin DIBO Alkyne to convert the azido-TMMs into biotin-TMMs. Surface-accessible biotin-TMMs were then detected by incubation with Alexa Fluor 488-conjugated streptavidin and visualized through fluorescence microscopy. Where indicated, compounds such as ethacridine and SQ109 were added 15 minutes prior to the addition of 6-azido-trehalose and maintained in all subsequent wash steps. Fluorescence readings were obtained using a BioTek Citation 5 multimode plate reader, with the detector set to an excitation wavelength of 488 nm and an emission wavelength of 525 nm. The percentage of inhibition was determined using the equation: % inhibition = 100 × (control − treated)/control. Fluorescence images were captured on an Olympus BX53 microscope at 100x magnification to visualize the distribution of labeled TMM on the spheroplast surface (Thacore & Willett, 1963)(Xu *et al*, 2017a).

### Measurement of mycobacterial transmembrane potential (ΔΨ)

We assessed the impact of ethacridine on the membrane potential of *Mtb* by employing a lipophilic cationic dye, 3,3-diethyloxacarbocyanine iodide (DiOC_2_). Following the method outlined in Chawla and Singh, 2013, mycobacterial cultures were labeled with DiOC_2_ for 15 minutes, subsequently washed with PBS (2–3 times), and resuspended to an OD_600_ of 0.2–0.3. Treatment was applied in triplicate to 500 μL of bacterial culture, with ethacridine and SQ109 administered at 1/2x, 1x, and 4x MIC concentrations.

Carbonyl cyanide m-chlorophenyl hydrazone (CCCP), a known protonophore, served as a positive control, while DMSO was included as a negative control. After 30 minutes, fluorescence intensity ratios of red to green (610/488 nm) were measured, where an increase in the red/green fluorescence ratio signified elevated membrane potential (Chawla & Singh, 2013) (Foss *et al*, 2016a).

### Flow Cytometry-Based Viability Analysis Using Sytox Green and Propidium Iodide

Flow cytometry-based analysis enables precisely discriminating live and dead bacterial cells by leveraging their distinct fluorescent properties. To assess the effect of drug treatment on Mycobacterium tuberculosis viability, flow cytometry was conducted using SYTOX Green (SG) and Propidium Iodide (PI), nucleic acid stains with distinct membrane permeability properties. SG and PI selectively enter cells with compromised membranes, enabling differentiation between live and dead bacterial populations. SG (Invitrogen, Cat. No. S7020) was diluted to a 20 μM working concentration in DMSO, while PI was prepared at a concentration optimized for bacterial staining. *Mtb* cultures in logarithmic growth phase were treated with the test drug ethacridine at 1x and 4x minimum inhibitory concentration (MIC), the reference drug SQ109 at 4x MIC, or DMSO (control). Untreated, unstained samples served as additional controls.

After drug exposure, bacterial cells were harvested, washed, and incubated with SG or PI for 15 minutes at room temperature in the dark. Stained samples were analyzed using a CyAn ADP flow cytometer (Beckman Coulter) equipped with a Cytek plate loader. Bacterial populations were gated based on forward and side scatter parameters to exclude debris. Fluorescent signals were used to quantify the proportion of cells with permeabilized membranes (non-viable cells). Histogram and dot plot analyses illustrated population shifts (Hendon-Dunn *et al*, 2016).

### In-silico studies

Molecular docking and molecular dynamics (MD) simulations of ethacridine were conducted to investigate its interaction with the Mmpl3 transporter protein, whose crystal structure (PDB ID: 6AJH) was retrieved from the Protein Data Bank (RCSB) (Zhang *et al*, 2019). The protein structure was co-crystallized with AU1235, which served as a reference for grid generation. Protein preparation was executed using Maestro software’s, Protein Preparation Wizard (PPW), with the OPLS2005 force field applied to minimize the structure. This preparation step included the removal of extraneous water molecules, non-essential ligands, and heteroatoms, followed by the addition of Kollman charges to ensure an optimized molecular setup. A docking grid was generated at the centroid of the co-crystallized AU1235 ligand, aligning the grid with the binding pocket to enhance docking accuracy. The 2D structure of ethacridine was sourced from PubChem and processed through Maestro’s LigPrep tool to generate multiple conformations and states. Molecular docking was performed using the Glide Dock extra-precision (XP) mode, to prioritize high-affinity binding modes (Friesner *et al*, 2006). and Following docking, MD simulations were executed in Maestro using the Desmond module, allowing for the observation of the protein-ligand complex in a dynamic aqueous environment (Bowers *et al*, 2006). This simulation offered insights into the stability and conformational shifts of ethacridine within the Mmpl3 binding site over time. This approach contribute valuable data on ethacridine’s potential as an Mmpl3 inhibitor (Stampolaki *et al*, 2023).

### Statistical Analysis

The statistical analysis was conducted using GraphPad Prism software, version 8. Non-linear regression analysis was applied to determine dose-response relationships, specifically employing a four-parameter logistic model to fit the log inhibitor concentration versus response curve. This approach allowed for the determination of key metrics, including the half-maximal inhibitory concentration (IC_50_) and slope, providing a robust model to evaluate each dataset. To compare the responses across different experimental groups, a non-parametric multiple comparison test (such as Dunn’s test or Kruskal-Wallis’s test) was used. This test identified statistically significant differences between groups with the following significance levels: *p* < 0.05 (*), *p* < 0.01 (**), *p* < 0.001 (***), and *p* < 0.0001 (****). Results meeting these criteria were considered significant.

## Author Contribution

N.P.K designed research; P.K.A. A.R., S.S, S.M., A.D., and A.S., performed research; P.K.A., and N.P.K analyzed data; P.K.A. and N.P.K. wrote the paper; and all authors contributed to writing the paper.

## Acknowledgement

We thank Prof. William R. Jacob, <u>Albert Einstein College of</u> <u>Medicine</u> for the gift of the derivative strains of *Mtb* H37Rv. This research is supported by the Department of Biotechnology, GOI, New Delhi, India under its (D.O. NO. BT/HRD/35/02/2006 (No. BT/RLF/66/2017)), and National Institute of Pharmaceutical Education and Research Hyderabad under the Department of Pharmaceuticals, Ministry of Chemicals and Fertilizers, Govt of India.

